# Integrins protect nociceptive neurons in models of paclitaxel-mediated peripheral sensory neuropathy

**DOI:** 10.1101/829655

**Authors:** Grace Ji-eun Shin, Maria Elena Pero, Luke A. Hammond, Anita Burgos, Samantha E. Galindo, Francesca Bartolini, Wesley B. Grueber

**Affiliations:** Columbia University, Zuckerman Mind Brain and Behavior Institute, Jerome L. Greene Science Center, 3227 Broadway, L9-007, New York, NY, 10027 USA; Columbia University, Pathology, New York, NY, 10032 USA; University of Naples Federico II, Naples, Italy; Columbia University, Neuroscience, New York, NY, 10032 USA

## Abstract

Chemotherapy induced peripheral neuropathy (CIPN) is a major side effect from cancer treatment with no known method for prevention or cure in clinics. CIPN primarily affects unmyelinated nociceptive sensory terminals. Despite the high prevalence of CIPN, molecular and cellular mechanisms that lead to CIPN are still poorly understood. Here, we used a genetically tractable *Drosophila* model and primary sensory neurons isolated from adult mouse to examine the mechanisms underlying CIPN and identify protective pathways. We found that chronic treatment of *Drosophila* larvae with paclitaxel caused sensory neuron degeneration, altered the terminal branching pattern of nociceptive neurons, and reduced thermal nociceptive responses. We found that nociceptive neuron-specific overexpression of integrins, which are known to support neuronal maintenance in several systems, conferred protection from paclitaxel-mediated cellular and behavioral phenotypes. Live imaging and superresolution approaches provide evidence that paclitaxel treatment causes cellular changes that are consistent with alterations in endosome-mediated trafficking of integrins. We used primary dorsal root ganglia neuron cultures to test conservation of integrin-mediated protection. We show that overexpression of a human integrin β subunit 1 (ITGB1) also prevented degeneration following paclitaxel treatment. Altogether, our study supports conserved mechanisms of paclitaxel-induced perturbation of integrin trafficking and a therapeutic potential of restoring integrin levels to antagonize paclitaxel-mediated toxicity in sensory neurons.

## Introduction

Chemotherapy-Induced Peripheral Neuropathy (CIPN) develops in over 60% of cancer patients and survivors (Seretny et al., 2014). CIPN significantly impacts quality of life as damage of sensory nerves may be permanent (Peltier and Russell, 2002), and is often a dose-limiting factor during cancer treatment. Patients with CIPN report pain-related symptoms including allodynia, hyper or hypoalgesia, or pain that can be more severe than the pain associated with the original cancer (Han and Smith, 2013). Despite increasing data on agents that protect sensory nerves, our limited understanding of the mechanisms of CIPN impedes effective treatment (Cavaletti and Marmiroli, 2010; Chua and Kroetz, 2017; Lisse et al., 2016). Studies from model systems may be helpful in identifying molecules that protect morphology from the effects of chemotherapeutics.

In the current study, we explored the mechanisms of CIPN induced by paclitaxel using two established models, *Drosophila* larval nociceptive neurons (Bhattacharya et al., 2012; Brazill et al., 2018) and primary DRG neurons isolated from adult mouse (Gornstein and Schwarz, 2017). Similar to other peripheral neuropathies, CIPN primarily affects unmyelinated intraepidermal nerve fibers that detect painful or noxious stimuli (Aley et al., 1996; Bennett et al., 2011; Lauria et al., 2003; Schmidt et al., 1997; Siau and Bennett, 2006; Tanner et al., 1998; Xiao et al., 2011). These small fibers are embedded in the epidermis, and continuously turn over coincident with the turnover of skin (Bennett et al., 2011). *Drosophila* class IV nociceptive neurons are a favored model for genetic studies of nociceptive neuron development and signaling mechanisms. Prior studies showed that class IV neuron morphology is sensitive to paclitaxel and demonstrated changes of nociceptive neurons at the onset and the end stage of CIPN (Bhattacharya et al., 2012; Brazill et al., 2018). Specifically, a chronic treatment at a high dose (30μM) induced fragmentation and dramatic simplification of branching of sensory terminals (Bhattacharya et al., 2012). Additionally, acute treatments of moderate doses (10- 20μM) induced hyperbranching of sensory arbors without changing the branch patterns (Brazill et al., 2018). Nociceptive neurons in *Drosophila* larvae detect multiple qualities of noxious stimuli, and project naked nerve terminals that are partially embedded in the epidermis (Hall and Treinin, 2011; Im and Galko, 2012). Larvae have a stereotyped behavioral response towards noxious stimuli that can serve as a readout of nociceptive neuron function (Burgos et al., 2018; Chen et al., 2016; Hwang et al., 2007; Poe et al., 2017). Nociceptive neurons in *Drosophila* larvae may therefore serve as a good *in vivo* model to study sensory changes induced by chemotherapeutics at morphological and functional levels.

Paclitaxel binds to tubulin and protect microtubules from disassembly. It is a commonly used chemotherapeutic drug for treatment of solid cancers such as breast, ovarian and lung cancers by virtue of its ability to inhibit cell division. Paclitaxel is more likely than other chemotherapeutics to cause chronic sensory neuropathy in patients and animal models (Gornstein and Schwarz, 2014; Hershman et al., 2011; Majithia et al., 2016; Shah et al., 2018; Wozniak et al., 2018; Xiao et al., 2011). Several CIPN animal and *in vitro* models have also revealed acute effects of paclitaxel (Brazill et al., 2018; Gornstein and Schwarz, 2017; Lisse et al., 2016; Pease-Raissi et al., 2017; Shemesh and Spira, 2010). While the mechanisms of acute and chronic neurodegeneration are likely to be distinct (Reichling and Levine, 2011), how long-term treatment of paclitaxel can affect sensory neuron morphology and function, and how neuronal arbors can be protected against long-term toxicity is not understood.

Several studies have shown that nociceptive sensory terminals share a close relationship with specific extracellular structures, most notably epidermal cells and extracellular matrix (ECM), yet whether paclitaxel impact this relationship is unknown. Integrins are a key mediator of the interaction between cells and the ECM, and impact neuronal maintenance in both vertebrate and invertebrate systems (Han et al., 2012; Kim et al., 2012; Moreno-Layseca et al., 2019; Nieuwenhuis et al., 2018). We therefore explored whether and how integrin-ECM interactions may impact sensory neuron maintenance upon paclitaxel-induced toxicity.

Here, we have used *Drosophila* and isolated mouse DRG neurons to investigate the pathological effect of paclitaxel in sensory neurons. Morphological changes occurred at paclitaxel doses that also caused changes in thermal nociceptive behaviors. Cell-specific overexpression of integrins protected nociceptive neurons from morphological alterations and prevented the thermal nociceptive behavior deficits caused by paclitaxel. This was consistent with the observation that paclitaxel affected the endosomal-lysosomal pathway and reduced integrin mediated recycling in *Drosophila*. Transduction of integrins also protected adult mouse DRG sensory neurons from paclitaxel-mediated toxicity *in vitro* indicating that integrin mediated protection is conserved in a vertebrate model of CIPN. Our study suggests that altered interactions between sensory neurons and their extracellular environment are an important contributor to paclitaxel mediated neuronal pathology.

## Results

### Paclitaxel alters the branching pattern of *Drosophila* nociceptive neurons

We first sought to confirm and extend prior results on the effect of paclitaxel on sensory dendrites using high-resolution analysis of terminal morphology. We administered paclitaxel (10, 20 and 30μM) in food beginning from 24-28h AEL (early 1^st^ instar). By this stage, da sensory neurons have completed axon pathfinding and formed major branches (Gao et al., 1999). All of these paclitaxel concentrations were previously used as models for CIPN (Bhattacharya et al., 2012; Brazill et al., 2018). We dissected treated larvae at the late 3^rd^ instar stage and found that while survival of larvae treated with 30μM was rare (<5% survival rate by 3^rd^ instar stages), animals tolerated 10 and 20μM well and often survived past 3^rd^ instar stages. da neuron branches still broadly covered their receptive field in larvae fed 10 and 20μM (Figure 1A-C), but neurons showed morphological changes such as pre-degenerative varicosities and degenerative fragmentation (Supplemental Figure 1, Figure 2). Nociceptive neurons in animals treated with 10μM did not show an overt change in branching pattern. By contrast, 20μM treatments induced marked changes in branching patterns. Peripheral arbors were sporadically clumped and patchy in proximal regions of the arbor and depleted in more distal regions (Figure 1A-C, Supplemental Figure 2A-D). Density analysis revealed that control nociceptive arbors have a peak at ∼5% (% of area that is occupied by branches within 50 *×* 50 μm^2^ (Figure 1D Supplemental Figure 2A). By contrast, paclitaxel dose-dependently increased overall local density of nociceptive arbors which showed no clear peaks in their distribution (Figure 1D). By plotting the data according to the distance from soma, we found that paclitaxel specifically caused increase in the proximal branch density (Figure 1E), also supported by Sholl analysis Supplemental Figure 1E). Neither concentration of paclitaxel caused a change in total dendrite length compared to control (Figure 1F). On the contrary, we found that 20μM, but not 10μM treatment increased the total number of dendrite branch points (Figure 1G). Thus, to explore the events that may precede degeneration of nociceptive arbors, we used both 10 and 20μM and focused on using 20μM of paclitaxel as a dose to further study the effects on *Drosophila* nociceptive arbors.

**Figure 1.**
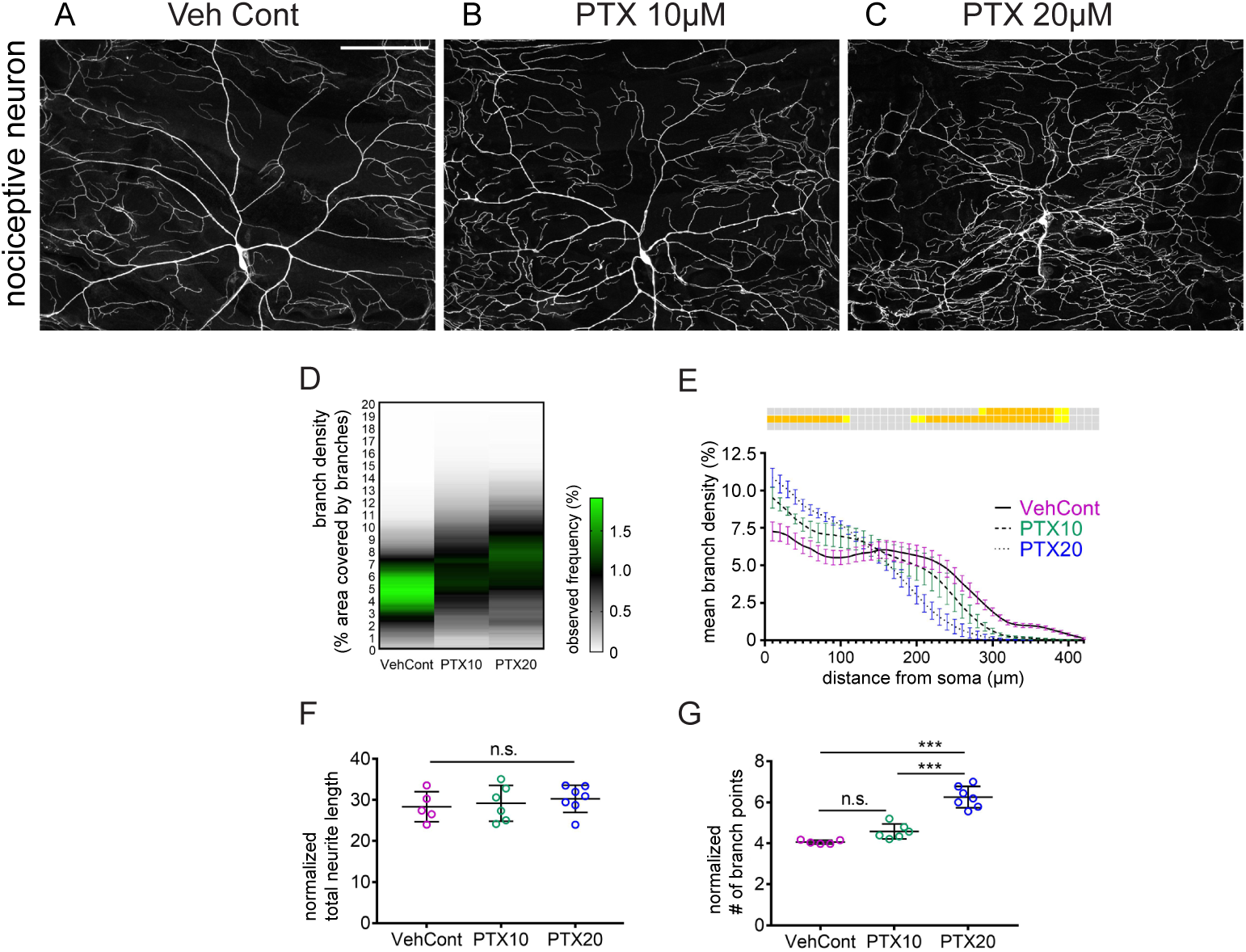
Paclitaxel causes changes in nociceptive neuron branch pattern in *Drosophila*. A-C. Micrographs from animals treated with vehicle control (A) and two different concentrations of paclitaxel [PTX; 10 (B) and 20μM (C)]. D, E. Density analyses of animals treated with vehicle control (N=7) and two different concentrations of paclitaxel (10 and 20μM, N=6 and 7, respectively). Branch density is calculated as described in Methods and Supplementary Figure 2. The same data are used in D and E except data in E are plotted according to distance from the neuronal soma. F, G. Quantification of nociceptive neuron morphology. (F) Total neurite length normalized against the area of the receptive field. (G) Neuron complexity measured by number of branch points normalized against total dendrite length. Scale bars=100μm. Error bars denote standard error of the mean (E) and standard deviation (F, G). Mixed effect analysis with Tukey’s multiple comparison post-test (E) Supplementary Table 1) and one-way ANOVA with Tukey’s multiple comparison post-test (F, G). Mean of each segment in density analysis (D) is color-coded, Statistical results from E are shown in color-coded boxes, *p<0.05 (yellow), **p<0.01 (orange), ***p<0.001 (red), n.s.>0.05 (gray).

**Figure 2.**
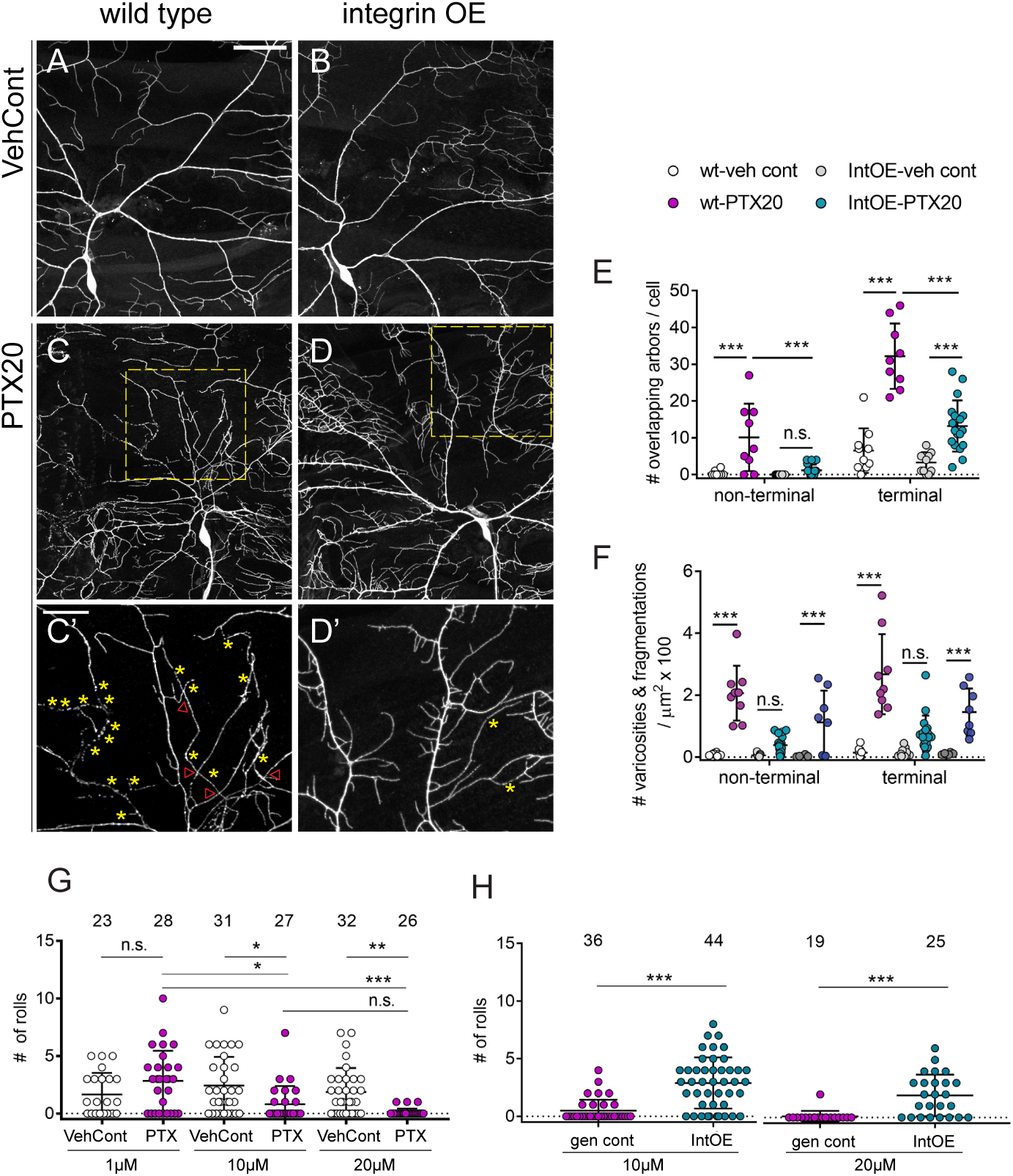
Cell-specific integrin overexpression prevents paclitaxel-induced morphological and functional changes in nociceptive neurons in *Drosophila*. A-D’. Micrographs of nociceptive neurons from wild type (A, C) and integrin overexpressing (B, D) animals treated with vehicle control or paclitaxel (20μM). C’, D’. Enlarged images of areas in C, D (yellow boxes) showing varicosities (yellow asterisks) and overlapping arbors (red arrow heads). See Supplementary Table 2 for a complete list of genotypes. G-H. Quantification of paclitaxel induced phenotypes, degeneration and overlapping arbors in non-terminal and terminal dendrites. Each data point refers to one larva represented by a single cell systemically selected from each larva. I-J. Quantification of nocifensive behavior evoked by global heat (40°C). Quantification of number of rolls within 60 seconds upon providing a thermal stimulus to the larva in control larvae (w^1118^) (I), genetic control (no UAS) and nociceptive neuron specific integrin overexpressing animals (J). Each data point represents a single animal. Scale bars=50μm (A-D) and 20μm (C’, D’). All error bars denote standard deviation. Two-way ANOVA with Tukey’s multiple comparison post-test (E, F), and Student’s t-test or Mann-Whitney test (G, H). *p<0.05, **p<0.01, ***p<0.001.

### Paclitaxel induced changes are prevented by cell-specific integrin overexpression in nociceptive sensory arbors

In addition to causing degeneration in nociceptive neurons, paclitaxel feeding resulted in extensive branch overlap, indicating a failure of self-avoidance (Figure 2A, C, C’, E, F). Two parallel mechanisms impact self-avoidance in *Drosophila* sensory dendrites. Dscam1-mediated recognition between sister dendrites results in repulsion and avoidance (Hughes et al., 2007; Matthews et al., 2007; Soba et al., 2007). In parallel, integrin receptors for the extracellular matrix maintain dendrites in an approximately 2D arrangement so dendrites come into contact with another rather than grow over or under each other (Grueber et al., 2002; Han et al., 2012; Kim et al., 2012). 3D crossing can be reduced or eliminated by overexpression of integrins. Overlapping arbors therefore raised the possibility that paclitaxel perturbs one of these self-avoidance mechanisms. Consistent with this hypothesis, overexpression of αPS1 and βPS integrins in nociceptive neurons significantly lessened paclitaxel-induced dendrite crossing (Figure 2B, D, D’, E). Remarkably, integrin overexpression also prevented paclitaxel-induced degenerative phenotypes (Figure 2D, D’, F). We compared this result with another cell adhesion receptor N-cadherin, that are also shown to promote cellular growth through cell-cell interaction (Doherty et al., 1991; Ferguson and Scherer, 2012). We found that N-cadherin overexpression in nociceptive neurons only partially prevented paclitaxel-mediated phenotypes (Supplemental Figure 3) arguing against the conclusion that integrin mediated protection is a mere consequence of a general increased adhesion of nociceptive neuron terminals (Supplemental Figure 3, compare with Figure 2E, F). Altogether, our results indicate that augmentation of integrins provides protection against paclitaxel-mediated neuropathy.

### Integrin overexpression prevents paclitaxel-induced change in thermal nociceptive response

*Drosophila* larvae show a stereotyped nocifensive behavior in response to thermal or mechanical noxious stimuli consisting of sequential C-shaped body bending, lateral rolling, and fast escape crawl (Burgos et al., 2018; Hwang et al., 2007; Ohyama et al., 2013; Tracey et al., 2003). We administered paclitaxel at 1, 10, and 20μM and tested for heat-induced nocifensive behavior. We observed a significant decrease in nocifensive responses at 10 and 20μM, but not at 1μM (Figure 2G). To determine if paclitaxel exerted an effect on sensory neurons or somewhere downstream of primary nociceptors, we performed a circuit epistasis experiment (Supplementary Figure 4). We fed larvae paclitaxel and activated downstream circuitry by expressing TrpA in Down and back (DnB) neurons, one of the main downstream targets of larval nociceptive neurons (Burgos et al., 2018). Thermogenetic activation of DnB neurons elicits bending and rolling behavior (Burgos et al., 2018), so we predicted that direct activation of DnB neurons would still be able to induce rolling behavior if the nociceptive neurons were the major target of paclitaxel. Indeed, activation of DnB neurons was as effective at inducing nociceptive behaviors with and without paclitaxel feeding. These results suggest that paclitaxel likely disrupts nociceptive behavior through action on primary sensory neurons rather than downstream circuitry.

We next examined whether integrin overexpression in nociceptive neurons can restore nocifensive behavior in paclitaxel-treated larvae. We found that changes in the levels of integrins or overexpression of N-cadherin did not alter nociceptive responses relative to genotype controls, although we did find that low and high integrin levels in nociceptive neurons lead to differences in the number of rolls initiated per nocifensive bout (Supplemental Figure 5). By contrast, at both 10 and 20μM of paclitaxel, overexpression of integrins in nociceptors led to a significantly stronger response to heat-induced rolling compared to genotype controls (Figure 2H). These results demonstrate that cell-specific integrin overexpression in nociceptive neurons prevents degeneration and disorganization of nociceptive neurons, and maintains function of these neurons in detecting noxious heat stimuli.

### Paclitaxel disrupts endosomal-lysosomal pathways and changes integrin trafficking in *Drosophila*

We next investigated the cellular changes in neurons caused by paclitaxel treatment that could be offset by augmenting integrin levels. Conceivably, these changes could include reduction in *de novo* integrin synthesis, reduced integrin recycling, and/or increased integrin degradation. Prior studies in other systems indicate that integrin levels are maintained by continuous recycling via tight regulation of the endosomal pathway rather than degradation and *de novo* synthesis (Moreno-Layseca et al., 2019). We therefore investigated potential changes caused by paclitaxel in endosomal and lysosomal pathways in nociceptive neurons in *Drosophila*. To initially assess overall changes in the endocytic pathway, we chose two markers, the small GTPase Rab4 and the transporter protein Spinster. Rab4 mediates the early endosome to recycling endosome transition and is found in both early and recycling endosomes (Hoogenraad et al., 2010; Sonnichsen et al., 2000; Vonderheit and Helenius, 2005). Spinster plays a critical role in the autophagy-lysosome transition and is found in late endosomes, lysosomes, and autophagosomes (Nakano et al., 2001; Rong et al., 2011). We expressed Rab4-RFP and Spinster-RFP in nociceptive neurons and monitored their dynamics in primary neurites in late third instar larvae.

High-resolution time-lapse images (see Methods) revealed a bidirectional trafficking of Rab4 positive endosomes of various sizes and speeds (Figure 3A). Unlike morphological changes, which showed paclitaxel dose-dependence, we found that the effects on Rab4 and Spinster trafficking were concentration independent (Figure 3C-G). Paclitaxel treatment caused a higher portion of Rab4 positive vesicles to become stationary at the expense of both anterograde and retrograde motility (Figure 3C) and the motile pool of Rab4 positive vesicles showed a reduced velocity and a higher frequency of direction switching (Figure 3D, E). Motile vesicles showed a significant reduction in density, whereas paclitaxel treatment did not affect the density of stationary vesicles (Figure 3F). In addition, both 10 and 20μM paclitaxel caused a marked enlargement of Spinster positive lysosomes. Together, our result demonstrates disruptions in both recycling endosome and lysosomal pathways by paclitaxel.

**Figure 3.**
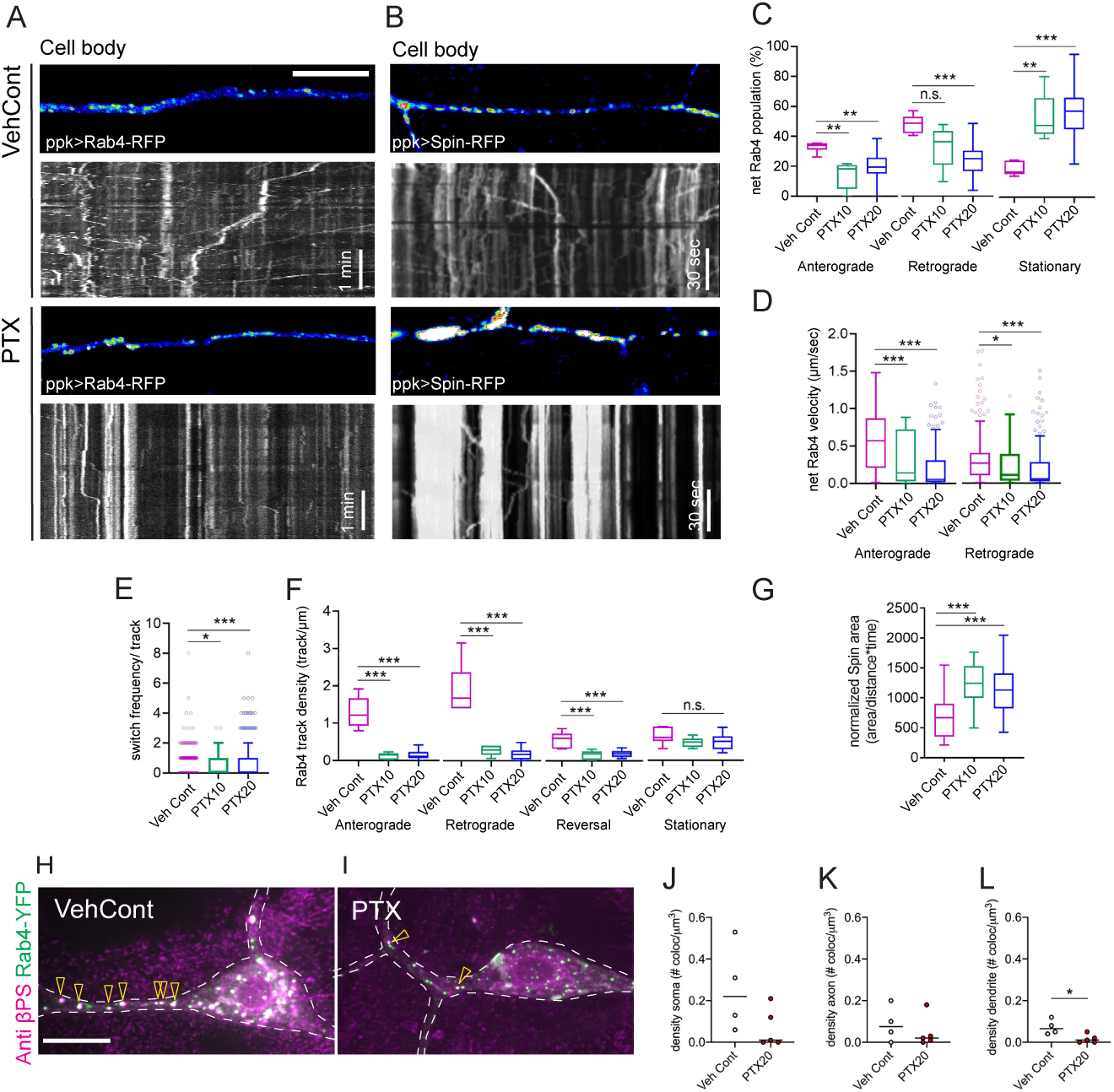
Paclitaxel disrupts endosomal-lysosomal pathways and changes integrin trafficking in *Drosophila*. A-B. Rab4-mRFP (recycling endosome marker, A) and Spinster-RFP (lysosome marker, B) were expressed in nociceptive neurons using nociceptive neuron specific driver ppk1.9Gal4. Rab4-RFP or Spinster-RFP are shown in nociceptor dendrite in still images. Movements are visualized kymographs in animals treated with vehicle and paclitaxel. For paclitaxel treated neurons, 10 and 20μM treatment resulted in qualitatively similar kymographs and only 20μM treated neurons are shown. C-F. Quantification of net Rab4 vesicle population showing anterograde or retrograde movement, or that are stationary (C, Vehicle control N=6, PTX10 N=5, PTX20 N=27, N denotes the number of individual kymographs), net velocity (D, Vehicle control n=247, PTX10 n=26, PTX20 n=167, n denotes the number of individual tracks), switch frequency (E, Vehicle control n=639, PTX10 n=91, PTX20 n=418, n denotes the number of individual tracks), and track density (F, Vehicle control N=6, PTX10 N=5, PTX20 N=27, N denotes the number of individual kymographs). G. Quantification of lysosomal compartment occupancy in dendrites. An area corresponding to 20 frames with minimal animal movement was selected from each kymograph. Fluorescent area under the curve was measured and normalized to distance selected for imaging. Each data point represents a kymograph collected from live imaging. One or two areas were selected from a neuron, and one or two neurons were selected in an animal. Vehicle control N=21, PTX10 N=25, PTX20 N=14, N denotes for the number of individual kymographs. H-L. Rab4-mRFP and integrins were co-expressed in nociceptive neurons by using nociceptive neuron specific driver ppk1.9Gal4 in animals treated with vehicle and paclitaxel (20μM). Soma, axon and dendrites (functionally nociceptive arbors) were segmented for quantification. H-I. Micrographs collected by using superresolution iSIM microscopy, visualizing integrin β subunit (βPS) by antibody labeling (endogenous + overexpressed) and Rab4 by antibody labelling against GFP (overexpressed only). Arrowheads denote co-localized puncta between integrin βPS and Rab4. J-L. Normalized density (per volume) of colocalized puncta between integrin βPS and Rab4 in soma (J), axon (K), and dendrite (L) Each data point refers to a single cell used for quantification. Scale bars=10μm. (C-L) All data were plotted using Tukey’s box and whiskers. Depending on normality, either one-way ANOVA with Tukey’s multiple comparison post-test or Kruskal-Wallis test with Dunn’s multiple comparison post-test was used. Welch’s t-test (J-L). *p<0.05, **p<0.01, ***p<0.001.

We next tested whether these changes could affect cargos such as integrins, that are required for the maintenance of nociceptive terminals. To demonstrate whether intracellular localization of integrins is changed due to paclitaxel mediated changes in endosomal-lysosomal pathways, we have expressed Rab4-YFP, *α*PS1 and *β*PS in nociceptive neurons (Figure 3H, I). We double labelled Rab4 and *β*PS integrins, and then imaged using superresolution microscopy (York et al., 2013) to quantify their co-localization in soma, dendrites and proximal axon. Consistent with the results obtained from live imaging of Rab4 trafficking, the number of Rab4 vesicles that colocalized with integrins in nociceptive arbors (dendrites) was significantly reduced by paclitaxel (Figure 3J-L). While we cannot exclude roles for degradation and *de novo* synthesis in the regulation of integrin levels, our data indicate that paclitaxel disrupts Rab4-mediated recycling of integrins. Together with the observed increase in lysosome compartment, these intracellular changes could affect the availability of integrins in support of nociceptive arbor maintenance.

### ITGB1 transduction in adult mouse DRG neurons prevents axon degeneration

We next asked whether integrin-mediated protection from paclitaxel is also observed in vertebrate neurons. Since augmentation of a single integrin subunit can recruit endogenous subunit partners (Condic, 2001; Huet-Calderwood et al., 2017), we chose to transduce the human integrin β1 subunit 1 (ITGB1), a common subunit in all heteromeric dimers in adult DRG neurons in rodents (Plantman et al., 2008; Tomaselli et al., 1993; Wallquist et al., 2004). Lentivirus mediated delivery of ITGB1 was carried out in adult DRG neurons at 5DIV prior to treatment with either vehicle or 50nM paclitaxel for 72 hours starting at 12DIV, an experimental paradigm that had been previously shown to induce early and late signs of axon degeneration (Gornstein and Schwarz, 2017). To better capture the changes in the cells in culture and to avoid selection bias, we have sampled large areas (2 x 2 mm^2^) for the quantification. We found that treating DRGs with paclitaxel resulted in axon loss and axon fragmentation (Figure 4A-B’) as shown by a significant decrease in total axon area and increased degeneration index (a ratio between fragmented axon and total axon areas) compared to control DRG neurons (Figure 4E, F, Supplemental Figure 5). Consistent with the protective capacity shown in *Drosophila* nociceptive neurons, DRG neurons transduced with ITGB1 prior to paclitaxel treatment showed no change in total axon area and a reduced degeneration index compared to paclitaxel treated control neurons (Figure 4E, F). These results demonstrate a conserved protective role for integrins in *Drosophila* and vertebrate CIPN models.

**Figure 4.**
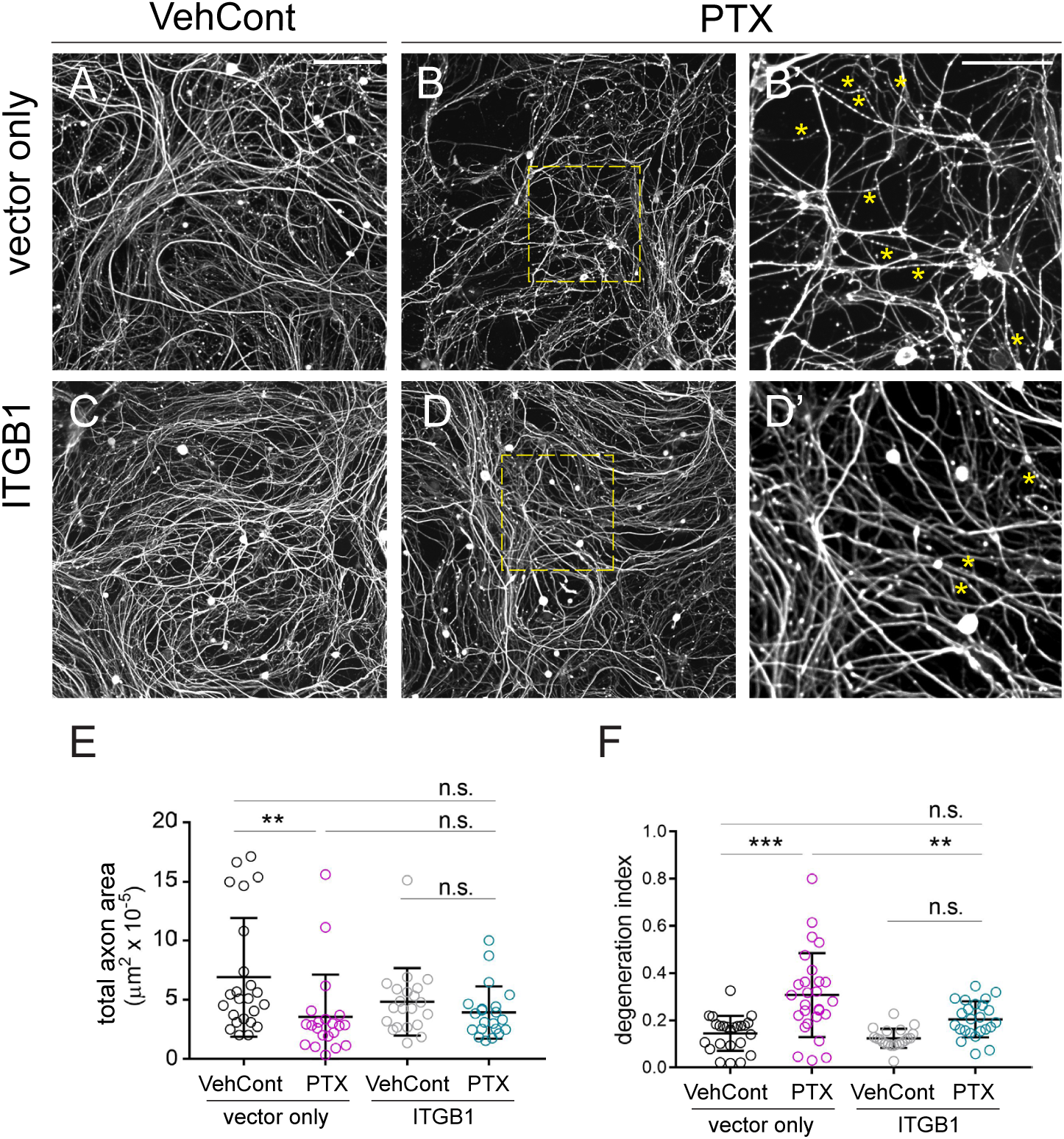
ITGB1 overexpression prevents paclitaxel-mediated degeneration in adult mouse DRG neurons *in vitro*. A-D. Micrographs DRG neurons treated with control virus and vehicle (A), control virus and paclitaxel (B), ITGB1 virus and vehicle (C), and control virus and paclitaxel (D). (B’, D’) Enlarged images of areas in B, D (dotted yellow boxes) showing fragmentation (yellow asterisks). E-F. Quantification of total area of axon (E) and axon degeneration (F) using degeneration index (degenerated area/total area of NF200 positive signal), calculated across the entire image. Two non-overlapping areas (2 x 2 mm^2^) were randomly selected from each well, and each data point refers to each region of interest (ROI). Scale bars=200μm (A-D) and 100μm (B’, D’). All error bars denote standard deviation. Kruskal-Wallis test with Dunn’s multiple comparison post-test (E) and one-way ANOVA with Tukey’s multiple comparison post-test (F). *p<0.05, **p<0.01, ***p<0.001. See also Supplementary Table 1 for a full list of statistical results.

## Discussion

In this study, we used *Drosophila* larval sensory neurons and adult mouse DRG neurons in culture to investigate the underlying mechanisms of CIPN induced by paclitaxel. By using cellular, genetic, and behavioral approaches, we demonstrate that integrin overexpression protects sensory neurons from paclitaxel treatment. We identified paclitaxel-mediated morphological changes in nociceptive neurons, and report quantitative changes in branch pattern arising from paclitaxel exposure. We report that overexpressing integrins, cell surface receptors critical during development and maintenance of non-neuronal cells and neurons (Hynes, 1992; Moreno-Layseca et al., 2019), protects nociceptive neuron morphology and function in *Drosophila*, and reduces axon degeneration in adult DRG neurons *in vitro*. We combined live imaging, high resolution and superresolution microscopy to identify the intracellular mechanisms that underlie integrin-mediated protection. Our results show that paclitaxel alters the endosomal-lysosomal pathway, and reduces recycling of integrins *in vivo*. Based on these data, we propose that paclitaxel alters intracellular pathways within nociceptive neurons and thereby disrupts the delivery of key cell surface receptors involved in interactions with the extracellular environment. This disruption leads to deficits in arbor maintenance, and eventually degeneration of nociceptive terminals.

### Integrins provide a link between sensory neurons and the extracellular environment

Intraepidermal nerve fibers (IENFs) are highly sensitive peripheral sensory structures because of their caliber and location adjacent to a highly dynamic epidermis (Bennett et al., 2011). Moreover, in the context of CIPN, recent studies in zebrafish indicate that epidermal cells are directly affected by paclitaxel and that epidermal changes precede neuronal degradation. In this system, blocking MMP13 signals that promote the degradation of the extracellular matrix and epidermis could partially rescue the neuronal damage caused by paclitaxel (Lisse et al., 2016). These prior studies indicate that degradation of neuronal substrates contributes to degeneration of adjacent arbors (Lisse et al., 2016). It is therefore essential to determine how sensory terminals are maintained in the context of a dynamic extracellular environment that can itself become sensitized to chemotherapeutic insults. Integrins are key cell surface receptors that link neuronal membranes to the extracellular matrix and our data show that they are a strong candidate factor that could mitigate the neuropathic effects of certain chemotherapeutics. Identification of the ligands that are involved in this protection is an important goal for future studies. Alternative signaling pathways could also contribute to integrin-mediated protection. For example, during the development of larval nociceptive neurons integrins also form a complex with the conserved receptor tyrosine kinase Ret (Soba et al., 2015). Together, they regulate the branching pattern of nociceptive neurons through epidermal derived TGF-beta Maverick. This complex is required for regular patterning, dynamic growth, and adhesion of dendritic branches (Hoyer et al., 2018; Soba et al., 2015). Since adhesive roles for the Ret-integrin complex are conserved in vertebrates (Cockburn et al., 2010), it is possible that similar co-receptors and ligands may interact with integrins to mediate interactions between IENFs and their extracellular environment.

### Evidence that endocytic changes caused by paclitaxel perturb integrin recycling

We provide evidence that paclitaxel disrupts trafficking of recycling endosomes in *Drosophila* peripheral nociceptive arbors. Specifically, we found a decrease in small GTPase Rab4 motility and abundance, and a decrease in integrin colocalization with Rab4 vesicles. Rab4 acts at the interface between early/sorting endosomes and recycling endosomes, and is a key mediator of integrin recycling (Arjonen et al., 2012; Moreno-Layseca et al., 2019; Roberts et al., 2001). Perturbation of Rab4 motility and availability would likely lead to reduced integrin turnover. By contrast, we found enlargement of Spinster positive lysosomal compartments in nociceptive neurons upon paclitaxel treatment. We have not directly examined whether this change impacts lysosomal activity or integrin degradation. Notably, prior studies in human non-small cell lung cancer cells *in vitro* have shown that paclitaxel upregulates the release and activation of lysosomal protease cathepsin B (Broker et al., 2004). Similarly, an increase in the activity of lysosomal enzymes was also reported in mouse hepatocytes *in vivo* following paclitaxel treatment (Krol, 1998). The exact mechanism leading to any of these changes is not known, however, the studies together suggest that paclitaxel commonly affects lysosomes in cancer cells, non-neuronal cells, and sensory neurons.

Interestingly, an opposing change of Rab4 and Spinster expression in our CIPN model also adds to a growing list of survival and maintenance factors that are modulated upon paclitaxel treatment. Paclitaxel reduces axonal expression of the pro-survival factor Bclw, but not Bcl2 or Bclx, in embryonic sensory neuron culture (Pease-Raissi et al., 2017). MMP13 was selectively activated in epidermal cells but not in neurons upon paclitaxel treatment (Lisse et al., 2016). Likewise, integrins may be one of several cargos that is reduced in the plasma membrane as a consequence of paclitaxel-mediated disruption in endo-lysosomal compartments. Integrin-mediated pathways may be more susceptible to these changes because integrin surface abundance relies heavily on the recycling and integrins may be continuously required in dynamic cellular compartments, such as nociceptive terminals or the leading edge of cancer cells (Arjonen et al., 2012; Han et al., 2012; Kim et al., 2012; Roberts et al., 2001; Roberts et al., 2004). Therefore, the integrin pathway may be relatively more strongly affected by changes in endosomal pathways.

### Substrate interactions in CIPN

Intraepidermal fiber (IENF) density correlates with severity of sensory peripheral neuropathy in patients, and the change in morphology serves as the strongest predictor of symptoms and prognosis in clinics (Hamid et al., 2014; Han and Smith, 2013; Khoshnoodi et al., 2016; Polydefkis et al., 2002; Tseng et al., 2006). In patients with chronic peripheral neuropathy, axon terminals are present in the subepidermal layer (Kennedy et al., 1996), but absent in the epidermis. It is therefore critical to understand the factors that help maintain terminal nerve fibers and that could prevent their degeneration in CIPN. Our results show that integrin supplementation promotes the maintenance of sensory terminals upon paclitaxel treatment, suggesting that integrins could be a key factor in somatosensory terminal maintenance to counteract CIPN. Our results suggest that integrin supplementation promotes the maintenance of sensory terminals upon paclitaxel treatment. Notably, other studies have shown that integrin levels are increased after neuron injury, and levels correlate with regenerative ability (Nieuwenhuis et al., 2018). Differences in integrin levels between peripheral DRG neurons and neurons in the CNS may correlate with different abilities of these neurons to regenerate (Andrews et al., 2016).

Currently, there is no effective method of preventing or treating CIPN other than stopping chemotherapeutic treatment or changing the chemotherapy regimen. More than 80% of CIPN patients treated with paclitaxel still report symptoms six months after the cessation of chemotherapeutic treatment (Hershman et al., 2011; Majithia et al., 2016). Limited symptomatic relief is provided by opioid analgesics, antidepressants, or anticonvulsants (Majithia et al., 2016; Shah et al., 2018). Our results provide an *in vivo* model to study CIPN-induced changes in neurons, and have identified and characterized integrins as a protective pathway. There is accumulating evidence from the literature and clinics that underscore the importance of intrinsic mechanisms in preventing neural degeneration. Our current study provides evidence for changes in the ability of neurons to link to, and interact with, the extracellular environment. Further studies of substrate interactions in CIPN models might provide important insights into the mechanisms, etiology, and treatment of this condition.

## Materials and methods

### Fly stocks

UAS-αPS1 (*multiple edematous wings, mew*) on III and UAS-βPS (*myospheroid, mys*) on II were provided by Dr. K. Broadie (Vanderbilt University). UAS-Ncad^7b-13a-18b^ (on II) line was provided by Dr. Chi-Hon Lee (National Institute of Health). Ppk1.9-Gal4 (II), ppkcd4tdgfp (III) and ppkcd4tdtom (III), and 412-Gal4 have been described previously (Ainsley et al., 2003; Burgos et al., 2018; Gohl et al., 2011; Han et al., 2011). W^1118^ and UAS-Rab4-mRFP, UAS-Spin-myc-mRFP, UAS-YFP-Rab4-WT, UAS-βPS-RNAi, UAS-TrpA1 lines were obtained from the Bloomington Stock Center (Bloomington, IN). Full genotypes of the animals used in the study can be found in Supplemental Table 2.

### Larval assay setup for paclitaxel treatment

Paclitaxel (Tocris Bioscience) was diluted to 5mM in ethanol, aliquoted and stored in −80°C until use. Fresh aliquots of paclitaxel were diluted to a final concentration of 1 – 30μM in 1 × PBS, and a matching concentration of ethanol was added as a vehicle control. This solution was used to make food using instant *Drosophila* medium (Formula 4-24 ®, Carolina Biological Supply Company) immediately before starting the larval assay. Embryos were collected on grape plates with yeast paste made with 0.5% propionic acid. First instar larvae (24 – 28h AEL) were collected manually with a paint brush, washed in 1 × PBS, and placed into paclitaxel-containing food. Larvae were reared at 25°C and were stage-matched to late third instar (based on the morphology of anterior spiracles) at the time of collection. Controls typically reached late third instar by 5-7 days. Any remaining animals that did not reach late third instar by 10 days after starting the treatment were discarded. All animals matching the developmental criteria were collected to avoid any selection bias.

### Global activation heat nocifensive assay and behavioral analysis

Global activation heat nocifensive assays were performed as previously described (Burgos et al., 2018) with a minor modification. Briefly, a single late third instar larva (stage-matched) was placed on a 0.6% black ink gel (Super Black India ink, Speedball) in 1% agarose in ddH_2_O heated to 40°C by a peltier device (CP-031, TE technology) equipped with a temperature controller (TC-36–25-RS232, TE technology). Animals with matching genotype to experimental groups except UAS transgenes were used as a genotype control. Where possible, paclitaxel treated animals were set up two-three days prior to control, so that both control and paclitaxel treated animals could be tested for nocifensive behavior on the same day. Behaviors were recorded for 60 seconds, and quantified blindly. The number of 360° rolls was scored by using trachea as a dorsal reference.

### Lentiviral packaging of human ecto-tagged ITGB1

Ecto-tagged human β1 integrin (ecto-GFP-ITGB1) in lentiviral expression vector (Huet-Calderwood et al., 2017) was kindly provided by David Calderwood’s lab at Yale University. pLENTI CMV Puro DEST (AddGene plamid #17452, gift from Eric Campeau) was purchased from AddGene. Lentiviruses containing pLENTI CMV-ecto-GFP-ITGB1 and a corresponding empty vector were produced according to Gornstein and Schwarz with some modification (Gornstein and Schwarz, 2017). Lentiviral expression vectors and packaging vectors (pLP1, pLP2, and pLP/VSVG, Thermofisher) were co-transfected to HEK293T cells using the calcium phosphate transfection method (Liu et al., 2014). Media containing viral particles were collected 24, 36, 48h after transfection, and lentiviral particles were filtered through 0.45μm, and then concentrated by centrifuging at 25,900 RPM at 4°C for 120 min (SW 32 Ti rotor, Beckman Coulter). The virus pellet was resuspended in Neurobasal media (Invitrogen) without any supplement to concentrate 250-fold from the packaging cell media, and stored in 30μl aliquots at −80°C until use. 5μl of lentivirus was added to 1ml culture at DIV 5, and ∼30% of the media was changed before paclitaxel administration at DIV 12. Expression of ITGB1 was checked by western blot.

### Adult mouse DRG culture and paclitaxel treatment

All protocols and procedures used in this study to prepare primary culture of DRG neurons were approved by the Committee on the Ethics of Animal Experiments of Columbia University and according to Guide for the Care and Use of Laboratory Animals distributed by the National Institutes of Health. Dorsal root ganglia from 8- to 12-week old C57Bl/6J mice of both sexes were dissected, dissociated and plated in a 12-well plate. Specifically, the spinal column was isolated immediately from a sacrificed mouse, and cut into approximately four pieces, including at the midline, to expose DRG for isolation. Each piece was dissected within 5 minutes, and the remaining tissue was placed on ice. Dissected DRG from cervical, thoracic and lumbar anatomical regions was immediately placed in HBSS (Life Technologies) or DMEM (Life Technologies) on ice. Once DRGs were collected, visible traces of axon tracts were cleaned before incubating in 1mg/mL Collagenase A (Sigma) for 1h at 37 °C. Following incubation, DRGs were washed twice in DMEM followed by 0.05% trypsin (Life Technologies) in DMEM digestion for 2 minutes at 37°C. After the digestion, cells were washed twice with Neurobasal medium supplemented with 2% B-27 (Invitrogen), 0.5mM glutamine (Invitrogen), 10% FBS (Sigma) and 100U/mL penicillin-streptomycin. DRG ganglia were then dissociated by repeated gentle pipetting until no clump was visible. Once dissociation was complete, cells were resuspended in supplemented Neurobasal media with 10% FBS to 1.8 ml (150μl x 12 wells) and plated onto 12 well plates (over 18mm coverslips) that had been coated overnight with 100μg/mL poly-D-lysine (Sigma) at 37°C and 10μg/mL laminin (Life Technologies) for 1h at 37 °C. After 30 minutes, 850μl of supplemented Neurobasal media was added to each well without disturbing plated cells. At 4DIV, ∼30% of the media was changed and 10μM AraC (Sigma) was administered to suppress growth of non-neuronal cells in culture. Approximately 30% of the media was again changed at 12DIV prior to paclitaxel treatment. 1μl of 1000 × paclitaxel (stock concentration 50μM) was added to each well containing 1ml of media to achieve a final concentration of 50nM. 1μl of DMSO was added as a vehicle control.

### Immunolabelling

Immunolabelling of *Drosophila* larvae was performed largely as described previously (Matthews et al., 2007). Briefly, late third instar larvae were dissected in 1 × PBS within 10 minutes to minimize degeneration during dissection. Dissected larvae were immediately fixed in 4% paraformaldehyde (PFA, Electron Microscopy Sciences) in 1 × PBS for 15 min, rinsed three times in 1 × PBS + 0.3% Triton X-100 (PBS-TX) for permeabilization, and blocked for 1 hr at room temperature in 5% normal donkey serum (NDS) in PBS-TX (Jackson Immunoresearch). Primary antibodies include chicken anti-GFP (1:1000; Abcam), rabbit anti-DsRed (1:250, Clontech), and mouse anti-mys (1:10, CF.6G11, Developmental Studies Hybridoma Bank), diluted in 5% NDS in PBS-TX. Primary antibodies were incubated overnight at 4°C, and washed in PBS-TX for 3 × 15 min. Species-specific, fluorophore-conjugated secondary antibodies (Jackson ImmunoResearch) were used at 1:1000 in 5% NDS in PBS-TX, and incubated overnight 4°C. Tissue was rinsed in PBS-TX for 3 × 15 min with a final rinse in PBS. Tissue was mounted on poly-L-lysine coated coverslips, dehydrated 5 minutes each in an ascending ethanol series (30, 50, 70, 95, 2 × 100%), cleared in xylenes (2 × 10 min), and mounted in DPX (Fluka).

For immunolabelling of DRG neurons, cultures were fixed in 4% PFA in PBS for 10 min, permeabilized with 0.1% Triton-X in PBS for 10 min, and blocked in 10% normal goat serum in PBS (NGS-PBS) for 1h. Primary antibodies were chicken NF200 (1:300, Aves Labs **Catalog # NFH**), rabbit ITGB1 (1:200, Invitrogen, **Catalog #** PA5-29606), which were diluted in NGS-PBS and incubated overnight at 4°C. Nuclei were stained with Hoeschst 33342 (1:1000, Sigma), which was incubated with primary antibodies. Species-specific, fluorophore-conjugated secondary antibodies (Invitrogen) were used at 1:500 in 10% NGS-PBS and incubated for 2h at the room temperature. Immunolabelled cells were then washed in 1 × PBS and mounted in Fluoromount-G (Southern Biotech).

### Image acquisition

*Drosophila* sensory neurons were imaged on a Nikon A1R laser scanning confocal microscope using a 60x 1.4 NA Plan Apochromat objective and a Yokogawa W1 spinning disk confocal microscope mounted on a Nikon Ti2-E stand using a 20x 0.75 NA Plan Apochromat objective. To avoid any selection bias, all animals from each experiment were imaged, and up to four ddaC nociceptive neurons were selected from abdominal segments. These cells were systematically and sequentially imaged in z-stacks from anterior to posterior positions in each larva. One cell per larva was systemically selected for quantification. For mouse DRG neurons Yokogawa W1 spinning disk confocal microscope was used. Images were collected initially in a 6mm × 6mm square in 2D, avoiding the peripheral region of the cultured cells. Three random points were selected from this image, which served as the center of non-overlapping z-stack images of 2mm × 2mm each.

### Image analysis

Prior to tracing of nociceptive neurons, an image containing a single neuron was blinded and the background was cleaned using background subtraction and non-local means denoising in ImageJ/Fiji. Because axons would be confounded with peripheral terminals during automatic detection, they were manually removed together with any neuronal arbors from neurons in adjacent segments. To improve a better automatic detection of arbors, brightness and contrast were optimized before processing in Vaa3D. The Vaa3D neuron 2 auto tracing plugin (Liu et al., 2018) was used to generate tracings of neuronal arbors. Any errors found in an automatically traced file were manually corrected via manual annotation using editing tools included in the neuron 2 plugin. The global neuron feature option was used to extract quantitative information from the reconstruction.

All data collected from *Drosophila* was randomized and blinded prior to quantification using customized python codes. Degeneration and overlap occurrences, were scored independently in terminal neurites and all other non-terminal neurites using the ImageJ/Fiji multi-point tool (Schindelin et al., 2012). We counted varicosities and fragmentation as early and late signs of degeneration, respectively. For quantifying branch overlap, we scored terminal dendrite overlap when at least one arbor was terminal neurite. All imaging data collected from DRG axons was blinded prior to quantification. For quantifying DRG degeneration phenotypes, a degeneration index was calculated as described in Gerdts et al., 2011 (Gerdts et al., 2011). Quantification was performed blindly using ImageJ/Fiji by an independent researcher who was not involved in experimentation and imaging. Briefly, images were automatically thresholded (global threshold) using a default auto-threshold method, binarized, and then total and fragmented axon areas were measured by using the particle analyzer plugin of ImageJ/Fiji. A circularity, determined as (area) / (*π* × radius^2^) and was calculated for each particle. Any particle of higher than 0.2 for circularity was considered fragmented. The degeneration index was calculated as a ratio of fragmented axon area over total axon area (Gerdts et al., 2011).

The local density of *Drosophila* sensory arbors was measured using a two-step custom ImageJ/Fiji macro. All images used for local density analysis were collected using identical settings on a Nikon A1R scanning laser confocal microscope. performing density analysis, the image files were pre-processed as described for the degeneration analysis. Local density was measured by a repeated measurement (sampled every 8 px in a 2048 x 2048 px image) of % of area occupied by the branches within a 50 x 50 μm^2^ square area, ignoring soma. Using these values a heatmap was generated which shows absolute levels of density (up to 15%) in assigned colors so that different images can be directly compared for their density levels and distribution. To generate local density relative to distance from the soma, the values from the heatmap were binned according to the distance from soma and mean and standard deviation was calculated for each neuron. To calculate the distribution of density values across the entire neuron, a custom Python script was used to measure the area of receptive field of each neuron, and the number of pixels in a given density (0-20%) in each neuron. For Sholl analysis, images containing a single neuron was prepared as described above and Sholl Analysis ImageJ/Fiji plugin was used at an interval of 5μm, starting from 10μm away from soma (Ferreira et al., 2014).

### Live imaging and quantitation of Rab4 and lysosomes in *Drosophila* larvae

Preparation of larvae for live imaging was largely performed as previously described (Chen et al., 2016). Briefly, a drop of fresh 3% agarose in ddH_2_O was placed on a slide and dried in the oven at 65°C overnight before use. A single larva was cleaned, placed on the dried agarose patch, and gently squeezed by a coverslip (size #1, Thorlab). This sandwich was held in place with sticky tape. Once mounted, the larva was imaged within 45 minutes. Live imaging was performed using Nikon A1R point-scanning laser confocal microscope (x40 objective, oil, 1.3 NA, 200μm WD, CFI60 Plan Fluor). Nociceptive neurons were identified by their expression of ppkCD4-tdGFP and a single image was collected as a reference map for the selection of ROIs. ROIs (256 x 128 px) were selected to image ∼50μm of primary dendrite at a Nyquist resolution. For both Rab4 and Spinster, we chose our ROIs at primary branches sufficiently away from soma, typically at around the first branch point (>100μm from soma). A single section of this ROI was imaged at 1.1sec/frame, while manually adjusting the focus when necessary to counter the minor movement of the animal. Images were collected for up to 5 minutes depending on the stability of the mounted larva. All images were randomized and blinded before any correction or quantification. Images were corrected for minor movements using the Stackreg plugin for ImageJ/Fiji. For quantification of Rab4, we used the Kymoanalyzer v.1 plugin for ImageJ/Fiji (Neumann et al., 2017) without any modification to default settings, except the threshold range for static vesicle was adjusted to avoid false positive detection of trafficking due to the movement of animal. Upon quantification, a custom R script was used to extract values from different measurements of Rab4 vesicles. The extracted values included net vesicle population distribution (stationary, anterograde, and retrograde), net velocity of vesicles in anterograde and retrograde directions, switch frequency, and track density. For quantification of Spin vesicles, 20-second portion of each kymograph was selected and the fluorescent intensity of this portion, representing the % of area occupied by lysosomes in the dendrite, was measured in ImageJ/Fiji using the line plot tool. All values above 3000 a.u. (over 65535 a.u., in 16 bit) were binned as a maximum value, and the values were used to plot a distribution curve. From here, area under the curve was calculated and normalized by the length of the dendrite selected in the ROI.

### Superresolution imaging and quantification of integrins and Rab4

Images were either collected by using a vt-iSIM (VisiTech International) (York et al., 2013) mounted on DMi8 Leica microscope using 93x 1.3 NA HC PL APO glycerol objective. Acquisitions included the cell body, axon and dendrites of ddaC neurons. These images were subsequently deconvolved using Microvolution (20 iterations), and were blinded prior to object analysis in Imaris (Oxford Instrument). Briefly, a channel for the nociceptive neuron (ppkcd4tdgfp) was segmented to use as a mask to isolate all neuronal puncta of integrins and Rab4. Smoothing (at 0.130 μm) and background subtraction (at 3 μm) was performed. All voxels with intensity greater than 482 a.u. (over 65535 a.u., 16 bit) in objects were filtered in, and objects smaller than 200 voxels were excluded. Channels for integrin *b*PS (anti-*b*PS) and Rab4-YFP, respectively, were segmented using background subtraction (at 0.5 μm). Intensity threshold was set to 400 as a default choice for most samples but 600 or 850 a.u. was also used depending on the fluorescent intensity of each sample. Objects were filtered to exclude those with fewer than 10 voxels. Detected puncta were compartmentalized into their locations in soma, dendrite, and axon via manual annotation of neuronal surface into these three categories. Co-localization of puncta expressing both *b*PS and Rab4 were detected by categorizing Rab4 positive objects into single (Rab4 alone) or co-positive (Rab4 and integrins) structure.

### Statistical analysis

Prism 8 was used for all statistical analysis and plotting data. Gaussian distribution was determined by using D’Agostino and Pearson or Shaprio-Wilk normality test. Depending on this result, either parametric or non-parametric test was selected for each analysis. For comparing three or more groups, a post-test was performed to correct for a multiple comparison to obtain an exact p-value. Detailed description of statistical tests performed for each experiment is included in figure legends. Statistical significance was determined at *p<0.05, **p<0.01, and ***p<0.001.

## Acknowledgements

We would like to thank Thompson Family Foundation Initiative (TFFI) at Columbia University for funding, and leadership and members of TFFI for feedback and helpful discussion. We thank present and past Grueber lab and Bartolini lab members for helpful discussion and the feedback and Clotilde Huet-Calderwood and David Calderwood (Yale University) for providing ITGB1 plasmids. We would also like to thank Atul Kumar for helping with mouse experiment preparation and helpful discussion, Tanguy Lucas, Siqian Feng, Susumu Antoku and Lily Qu for cloning advice, Daniel Iascone for help with using Vaa3D plugin, Terry Hafer for help with live imaging preparation for larvae, Tommy Kahn and Robert Cudmore (Johns Hopkins) for Python codes for blinding files. We also would like thank Zuckerman Institute’s Cellular Imaging platform for the access to the microscopes.

## Author contribution

G.J.S. and W.B.G. conceived and designed the research, discussed results and their interpretation, and prepared the manuscript. F.B. closely advised on the mouse experiments, discussed the results and their interpretation, and critically read the manuscript. G.J.S. performed all the experiments, with the exception of behavior experiments which were performed and quantified by A.B. and S.E.G. M.E.P. trained G.J.S. in mouse experiments, and helped with preparation of DRG culture. G.J.S. and M.E.P. performed data analyses for mouse experiments, and G.J.S. and L.A.H. performed density analysis and co-localization analysis. All other data analyses were performed by G.J.S. L.A.H. developed density analysis method and wrote custom ImageJ/Fiji macro, R and Python code for the analysis. L.A.H. closely advised on all imaging parameters. G.J.S. performed all statistical analyses.

## Supplementary figures

**Supplementary Figure 1.**
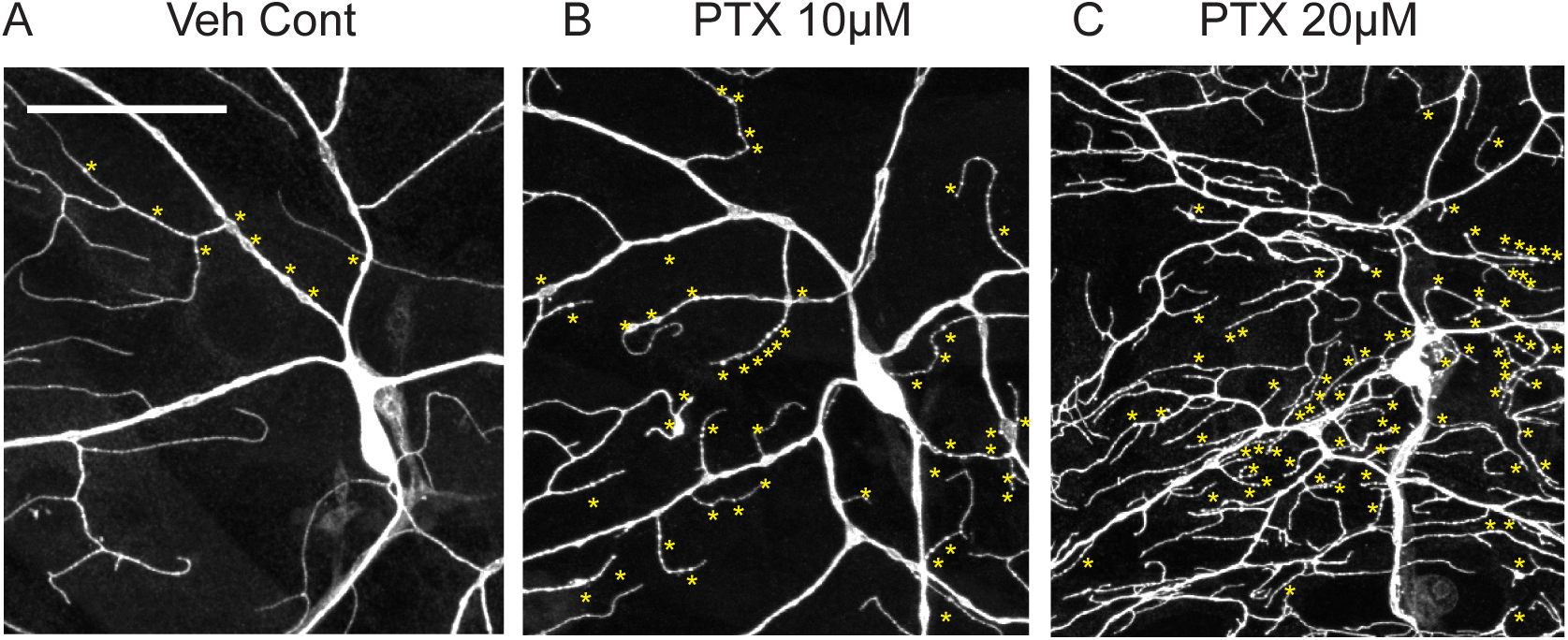
Degeneration phenotypes in neurons presented in Figure 1. Center of neurons shown in Figure 1A-C are enlarged to show signs of degeneration (varicosities and fragmentation, yellow asterisks). See also Figure 2. Scale bar = 50μm

**Supplementary Figure 2.**
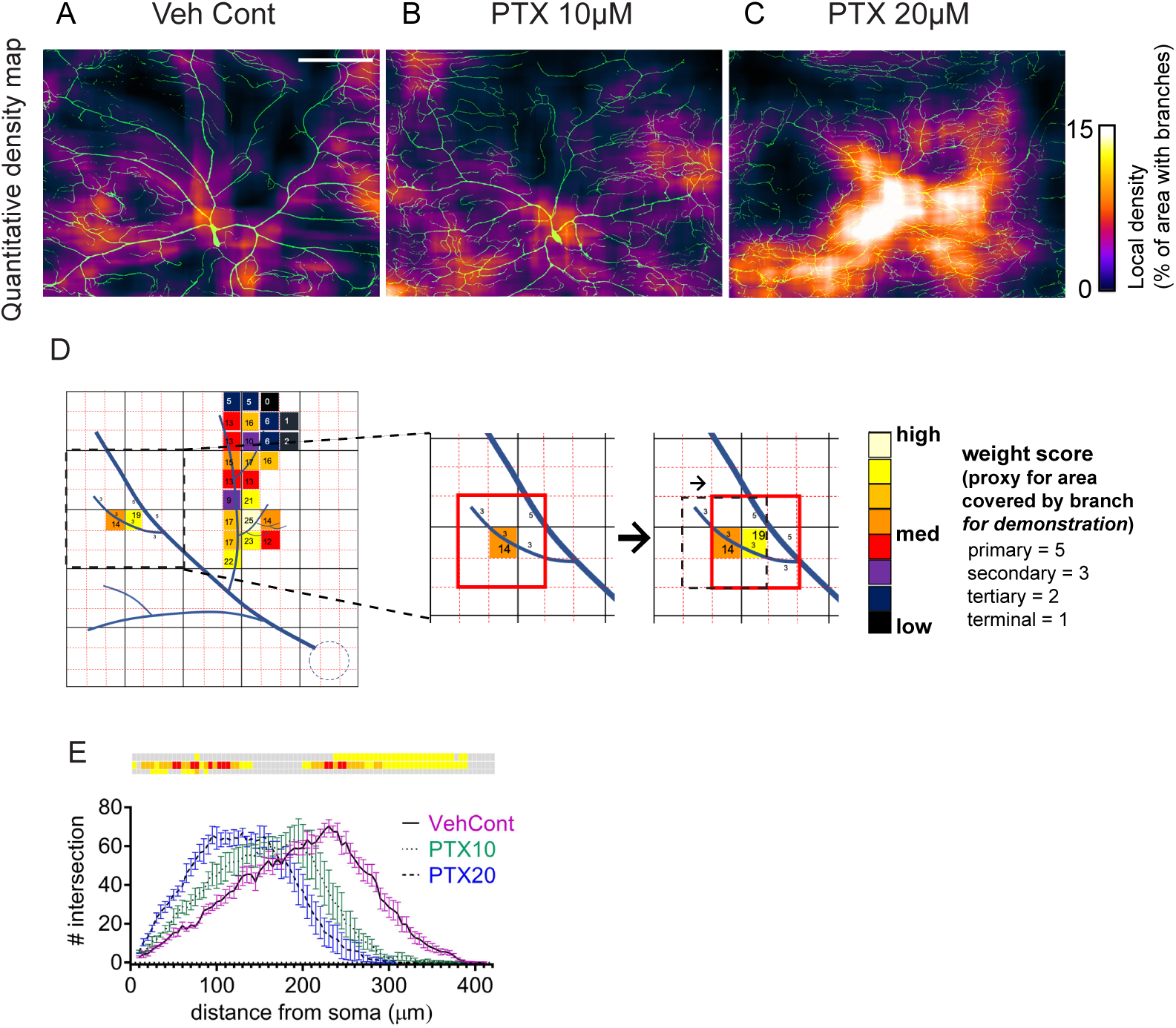
Quantitative visualization and the measurement of density by paclitaxel treatment. A-D. Examples of quantitative density heatmaps (A-C), and schematics illustrating the method of density analysis (D). A-C. To obtain heatmaps, the density (% area covered by branches in surrounding area sized at 50 x 50μm^2^) was measured every 2.5 μm (8 px in 2048 x 2048 image, also refer to method), resulted in 65536 measurement points (256 x 256) per image obtained in our study. Most of the density values were lower than 15% (Figure 1D), therefore we assigned the color for each measurement point corresponding to the scale of 0-15% to create a heatmap. D. Simplified illustration of the density analysis. Here, we used weight score as a proxy for % of area covered by branches. Dotted red boxes indicate each measurement point (15 x 15 measurement points in the first schematic), and the red boxes refer to surrounding area (3 x 3 boxes) for a measurement point in the center. Each color-coded measurement point (dotted red boxes) shows the density of a respective surrounding area, calculated according to the weight score and assigned for a color that matches with the scale (high-med-low). Simulation of the density measurement of two measurement points are illustrated in magnified images. Weight score of a surrounding area (red box, 3 x 3 boxes) for the first and the second measurement points have density scores of 14 and 19. They are mid and mid-high densities for this schematic neuron, therefore had been assigned with dark orange and yellow colors in the scale. The red box moves along the entire image to calculate density for each measurement point to create a heatmap for an entire neuron. E. Sholl analysis of the same set of neurons used in density analysis (Figure 1D, E) Scale bar=100μm. Error bars denote standard error means. Mixed effect analysis with Tukey’s multiple comparison post-test (See also Supplementary Table 1). Statistical results from E are shown in color-coded boxes, *p<0.05 (yellow), **p<0.01 (orange), ***p<0.001 (red), n.s.>0.05 (gray).

**Supplementary Figure 3.**
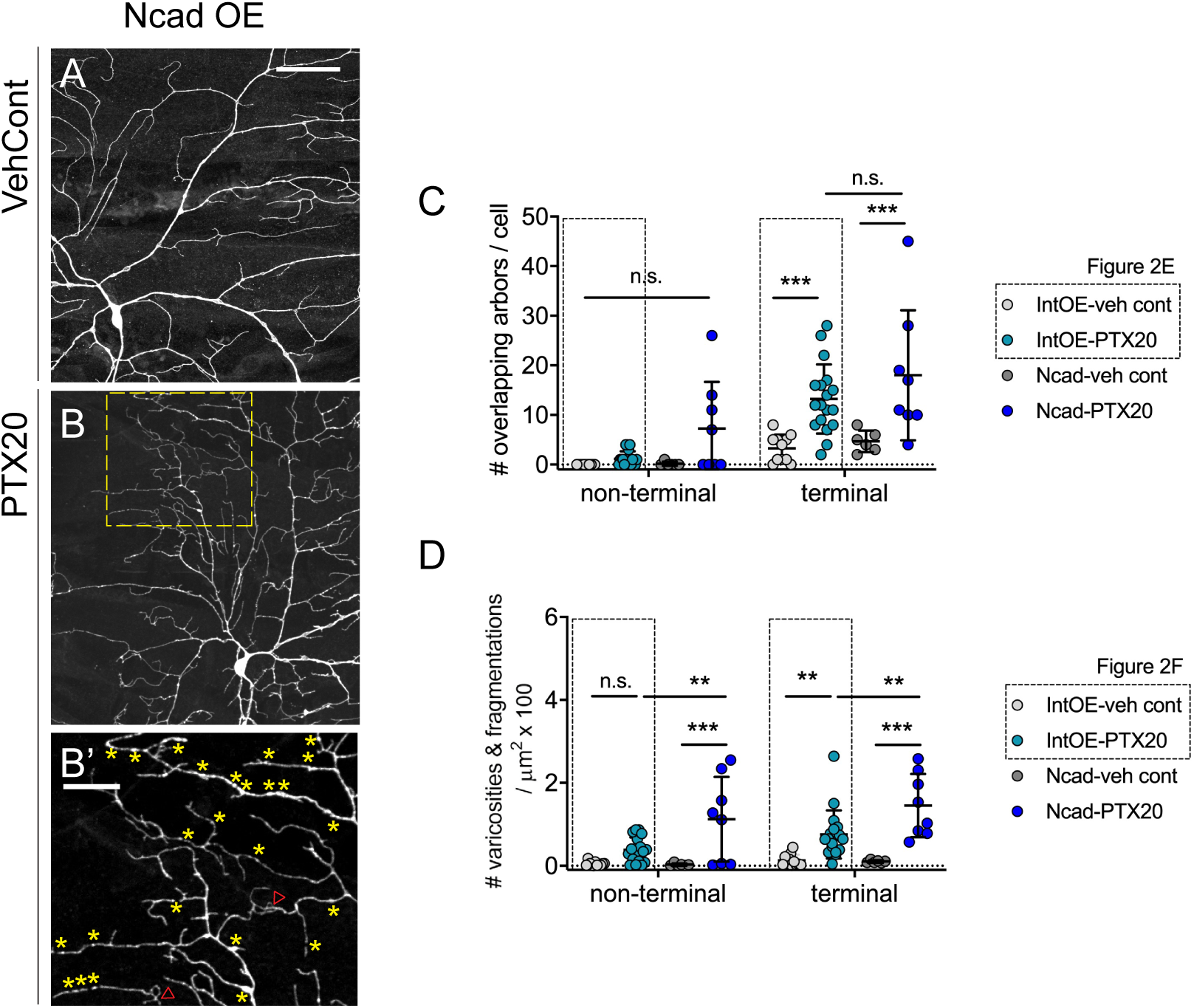
N-cadherin overexpression is partially protective against paclitaxel mediated morphological phenotypes. A-B’. Micrographs of nociceptive neurons from N-cadherin overexpressing animals treated with vehicle control and paclitaxel (20μM). B’. Enlarged images of areas in B (dotted yellow boxes) showing varicosities (yellow asterisks) and overlapping arbors (red arrow heads). C-D. Quantification of paclitaxel induced phenotypes, degeneration and overlapping arbors in non-terminal and terminal dendrites. Each data point refers to one larva represented by a single cell systemically selected from each larva. Scale bars=50μm (A, B) and 20μm (B’). All error bars denote standard deviation. Two-way ANOVA with Tukey’s multiple comparison post-test (C, D). *p<0.05, **p<0.01, ***p<0.001.

**Supplementary Figure 4.**
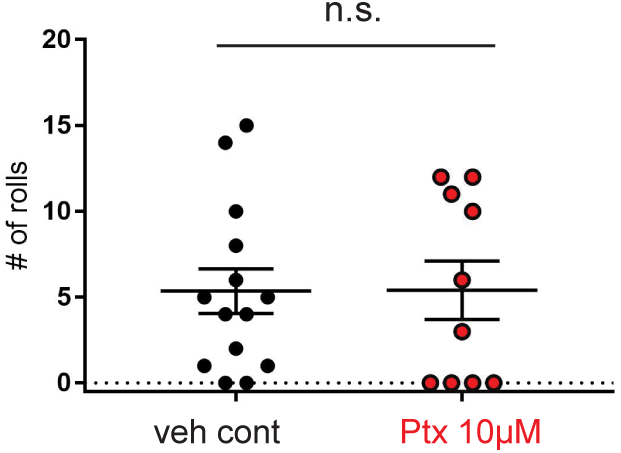
Nocifensive responses arising from thermogenetic activation of DnB interneurons is not sensitive to paclitaxel treatment. Animals expressing UAS TrpA under the control of 412Gal4 were reared until the wandering third instar stage. Animals were tested for rolling behavior induced by thermogenetic activation of DnB neurons. Quantification of rolling behavior was performed by counting the number of rolls within 60 seconds upon thermal stimuli to the larva. Mann-Whitney test. *p<0.05.

**Supplementary Figure 5.**
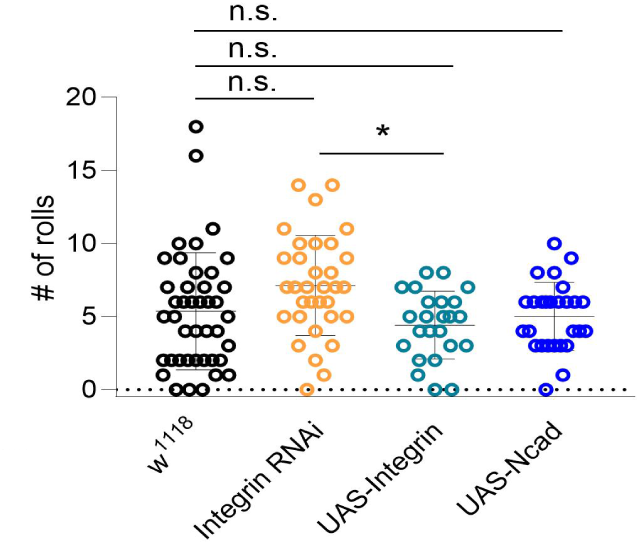
Neither changes in integrin levels nor N-cadherin overexpression alter nocifensive responses compared to control *Drosophila* larvae. Larvae of matching genotype to morphological analysis in Figure 2 were used for the baseline behavior analysis. Animals were reared until they reach wandering third instar stage and tested for nocifensive response against global thermal noxious stimuli (peltier device at 41°C). Quantification of rolling behavior was performed by counting the number of rolls within 60 seconds upon thermal stimulation of the larva. *p<0.05, Kruskal-Wallis test with Dunn’s multiple comparison post-test.

**Supplementary Figure 6.**
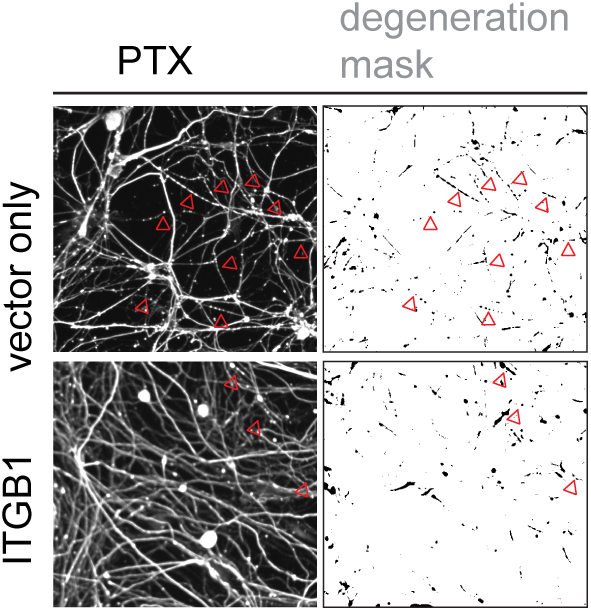
Examples of degeneration mask for calculating degeneration index. Degeneration mask was generated according to Gerdts et al., 2011. Arrow heads point to several examples of degenerating axons in the micrograph compared to matching locations within degeneration masks.

## Reference

Ainsley, J.A., Pettus, J.M., Bosenko, D., Gerstein, C.E., Zinkevich, N., Anderson, M.G., Adams, C.M., Welsh, M.J., and Johnson, W.A. (2003). Enhanced locomotion caused by loss of the Drosophila DEG/ENaC protein Pickpocket1. Curr Biol 13, 1557–1563.

Aley, K.O., Reichling, D.B., and Levine, J.D. (1996). Vincristine hyperalgesia in the rat: A model of painful vincristine neuropathy in humans. Neuroscience 73, 259–265.

Andrews, M.R., Soleman, S., Cheah, M., Tumbarello, D.A., Mason, M.R., Moloney, E., Verhaagen, J., Bensadoun, J.C., Schneider, B., Aebischer, P., et al. (2016). Axonal Localization of Integrins in the CNS Is Neuronal Type and Age Dependent. eNeuro 3.

Arjonen, A., Alanko, J., Veltel, S., and Ivaska, J. (2012). Distinct recycling of active and inactive beta1 integrins. Traffic 13, 610–625.

Bennett, G.J., Liu, G.K., Xiao, W.H., Jin, H.W., and Siau, C. (2011). Terminal arbor degeneration - a novel lesion produced by the antineoplastic agent paclitaxel. European Journal of Neuroscience 33, 1667–1676.

Bhattacharya, M.R., Gerdts, J., Naylor, S.A., Royse, E.X., Ebstein, S.Y., Sasaki, Y., Milbrandt, J., and DiAntonio, A. (2012). A model of toxic neuropathy in Drosophila reveals a role for MORN4 in promoting axonal degeneration. J Neurosci 32, 5054–5061.

Brazill, J.M., Cruz, B., Zhu, Y., and Zhai, R.G. (2018). Nmnat mitigates sensory dysfunction in a Drosophila model of paclitaxel-induced peripheral neuropathy. Dis Model Mech 11.

Broker, L.E., Huisman, C., Span, S.W., Rodriguez, J.A., Kruyt, F.A., and Giaccone, G. (2004). Cathepsin B mediates caspase-independent cell death induced by microtubule stabilizing agents in non-small cell lung cancer cells. Cancer Res 64, 27–30.

Burgos, A., Honjo, K., Ohyama, T., Qian, C.S., Shin, G.J., Gohl, D.M., Silies, M., Tracey, W.D., Zlatic, M., Cardona, A., et al. (2018). Nociceptive interneurons control modular motor pathways to promote escape behavior in Drosophila. Elife 7.

Cavaletti, G., and Marmiroli, P. (2010). Chemotherapy-induced peripheral neurotoxicity. Nat Rev Neurol 6, 657–666.

Chen, L., Nye, D.M., Stone, M.C., Weiner, A.T., Gheres, K.W., Xiong, X., Collins, C.A., and Rolls, M.M. (2016). Mitochondria and Caspases Tune Nmnat-Mediated Stabilization to Promote Axon Regeneration. PLoS Genet 12, e1006503.

Chua, K.C., and Kroetz, D.L. (2017). Genetic advances uncover mechanisms of chemotherapy-induced peripheral neuropathy. Clin Pharmacol Ther 101, 450–452.

Cockburn, J.G., Richardson, D.S., Gujral, T.S., and Mulligan, L.M. (2010). RET-mediated cell adhesion and migration require multiple integrin subunits. J Clin Endocrinol Metab 95, E342–346.

Condic, M.L. (2001). Adult neuronal regeneration induced by transgenic integrin expression. J Neurosci 21, 4782–4788.

Doherty, P., Rowett, L.H., Moore, S.E., Mann, D.A., and Walsh, F.S. (1991). Neurite outgrowth in response to transfected N-CAM and N-cadherin reveals fundamental differences in neuronal responsiveness to CAMs. Neuron 6, 247–258.

Ferguson, T.A., and Scherer, S.S. (2012). Neuronal cadherin (NCAD) increases sensory neurite formation and outgrowth on astrocytes. Neurosci Lett 522, 108–112.

Ferreira, T.A., Blackman, A.V., Oyrer, J., Jayabal, S., Chung, A.J., Watt, A.J., Sjostrom, P.J., and van Meyel, D.J. (2014). Neuronal morphometry directly from bitmap images. Nat Methods 11, 982–984.

Gao, F.B., Brenman, J.E., Jan, L.Y., and Jan, Y.N. (1999). Genes regulating dendritic outgrowth, branching, and routing in Drosophila. Genes Dev 13, 2549–2561.

Gerdts, J., Sasaki, Y., Vohra, B., Marasa, J., and Milbrandt, J. (2011). Image-based Screening Identifies Novel Roles for I kappa B Kinase and Glycogen Synthase Kinase 3 in Axonal Degeneration. Journal of Biological Chemistry 286, 28011–28018.

Gohl, D.M., Silies, M.A., Gao, X.J., Bhalerao, S., Luongo, F.J., Lin, C.C., Potter, C.J., and Clandinin, T.R. (2011). A versatile in vivo system for directed dissection of gene expression patterns. Nat Methods 8, 231–237.

Gornstein, E., and Schwarz, T.L. (2014). The paradox of paclitaxel neurotoxicity: Mechanisms and unanswered questions. Neuropharmacology 76 Pt A, 175–183.

Gornstein, E.L., and Schwarz, T.L. (2017). Neurotoxic mechanisms of paclitaxel are local to the distal axon and independent of transport defects. Exp Neurol 288, 153–166.

Grueber, W.B., Jan, L.Y., and Jan, Y.N. (2002). Tiling of the Drosophila epidermis by multidendritic sensory neurons. Development 129, 2867–2878.

Hall, D.H., and Treinin, M. (2011). How does morphology relate to function in sensory arbors? Trends Neurosci 34, 443–451.

Hamid, H.S., Mervak, C.M., Munch, A.E., Robell, N.J., Hayes, J.M., Porzio, M.T., Singleton, J.R., Smith, A.G., Feldman, E.L., and Lentz, S.I. (2014). Hyperglycemia- and neuropathy-induced changes in mitochondria within sensory nerves. Ann Clin Transl Neurol 1, 799–812.

Han, C., Jan, L.Y., and Jan, Y.N. (2011). Enhancer-driven membrane markers for analysis of nonautonomous mechanisms reveal neuron-glia interactions in Drosophila. Proc Natl Acad Sci U S A 108, 9673–9678.

Han, C., Wang, D., Soba, P., Zhu, S., Lin, X., Jan, L.Y., and Jan, Y.N. (2012). Integrins regulate repulsion-mediated dendritic patterning of drosophila sensory neurons by restricting dendrites in a 2D space. Neuron 73, 64–78.

Han, Y., and Smith, M.T. (2013). Pathobiology of cancer chemotherapy-induced peripheral neuropathy (CIPN). Front Pharmacol 4, 156.

Hershman, D.L., Weimer, L.H., Wang, A., Kranwinkel, G., Brafman, L., Fuentes, D., Awad, D., and Crew, K.D. (2011). Association between patient reported outcomes and quantitative sensory tests for measuring long-term neurotoxicity in breast cancer survivors treated with adjuvant paclitaxel chemotherapy. Breast Cancer Res Treat 125, 767–774.

Hoogenraad, C.C., Popa, I., Futai, K., Martinez-Sanchez, E., Wulf, P.S., van Vlijmen, T., Dortland, B.R., Oorschot, V., Govers, R., Monti, M., et al. (2010). Neuron specific Rab4 effector GRASP-1 coordinates membrane specialization and maturation of recycling endosomes. PLoS Biol 8, e1000283.

Hoyer, N., Zielke, P., Hu, C., Petersen, M., Sauter, K., Scharrenberg, R., Peng, Y., Kim, C.C., Han, C., Parrish, J.Z., et al. (2018). Ret and Substrate-Derived TGF-beta Maverick Regulate Space-Filling Dendrite Growth in Drosophila Sensory Neurons. Cell Rep 24, 2261–2272 e2265.

Huet-Calderwood, C., Rivera-Molina, F., Iwamoto, D.V., Kromann, E.B., Toomre, D., and Calderwood, D.A. (2017). Novel ecto-tagged integrins reveal their trafficking in live cells. Nat Commun 8, 570.

Hughes, M.E., Bortnick, R., Tsubouchi, A., Baumer, P., Kondo, M., Uemura, T., and Schmucker, D. (2007). Homophilic Dscam interactions control complex dendrite morphogenesis. Neuron 54, 417–427.

Hwang, R.Y., Zhong, L., Xu, Y., Johnson, T., Zhang, F., Deisseroth, K., and Tracey, W.D. (2007). Nociceptive neurons protect Drosophila larvae from parasitoid wasps. Curr Biol 17, 2105–2116.

Hynes, R.O. (1992). Integrins: versatility, modulation, and signaling in cell adhesion. Cell 69, 11–25.

Im, S.H., and Galko, M.J. (2012). Pokes, sunburn, and hot sauce: Drosophila as an emerging model for the biology of nociception. Dev Dyn 241, 16–26.

Kennedy, W.R., Wendelschafer-Crabb, G., and Johnson, T. (1996). Quantitation of epidermal nerves in diabetic neuropathy. Neurology 47, 1042–1048.

Khoshnoodi, M.A., Truelove, S., Burakgazi, A., Hoke, A., Mammen, A.L., and Polydefkis, M. (2016). Longitudinal Assessment of Small Fiber Neuropathy: Evidence of a Non-Length-Dependent Distal Axonopathy. JAMA Neurol 73, 684–690.

Kim, M.E., Shrestha, B.R., Blazeski, R., Mason, C.A., and Grueber, W.B. (2012). Integrins establish dendrite-substrate relationships that promote dendritic self-avoidance and patterning in drosophila sensory neurons. Neuron 73, 79–91.

Krol, T. (1998). Activity of lysosomal system in mouse liver after taxol administration. Gen Pharmacol 30, 239–243.

Lauria, G., Morbin, M., Lombardi, R., Borgna, M., Mazzoleni, G., Sghirlanzoni, A., and Pareyson, D. (2003). Axonal swellings predict the degeneration of epidermal nerve fibers in painful neuropathies. Neurology 61, 631–636.

Lisse, T.S., Middleton, L.J., Pellegrini, A.D., Martin, P.B., Spaulding, E.L., Lopes, O., Brochu, E.A., Carter, E.V., Waldron, A., and Rieger, S. (2016). Paclitaxel-induced epithelial damage and ectopic MMP-13 expression promotes neurotoxicity in zebrafish. Proc Natl Acad Sci U S A 113, E2189–2198.

Liu, J., Pasini, S., Shelanski, M.L., and Greene, L.A. (2014). Activating transcription factor 4 (ATF4) modulates post-synaptic development and dendritic spine morphology. Front Cell Neurosci 8, 177.

Liu, S.Q., Zhang, D.H., Song, Y., Peng, H.C., and Cai, W.D. (2018). Automated 3-D Neuron Tracing With Precise Branch Erasing and Confidence Controlled Back Tracking. Ieee T Med Imaging 37, 2441–2452.

Majithia, N., Temkin, S.M., Ruddy, K.J., Beutler, A.S., Hershman, D.L., and Loprinzi, C.L. (2016). National Cancer Institute-supported chemotherapy-induced peripheral neuropathy trials: outcomes and lessons. Support Care Cancer 24, 1439–1447.

Matthews, B.J., Kim, M.E., Flanagan, J.J., Hattori, D., Clemens, J.C., Zipursky, S.L., and Grueber, W.B. (2007). Dendrite self-avoidance is controlled by Dscam. Cell 129, 593–604.

Moreno-Layseca, P., Icha, J., Hamidi, H., and Ivaska, J. (2019). Integrin trafficking in cells and tissues. Nat Cell Biol.

Nakano, Y., Fujitani, K., Kurihara, J., Ragan, J., Usui-Aoki, K., Shimoda, L., Lukacsovich, T., Suzuki, K., Sezaki, M., Sano, Y., et al. (2001). Mutations in the novel membrane protein spinster interfere with programmed cell death and cause neural degeneration in Drosophila melanogaster. Mol Cell Biol 21, 3775–3788.

Neumann, S., Chassefeyre, R., Campbell, G.E., and Encalada, S.E. (2017). KymoAnalyzer: a software tool for the quantitative analysis of intracellular transport in neurons. Traffic 18, 71–88.

Nieuwenhuis, B., Haenzi, B., Andrews, M.R., Verhaagen, J., and Fawcett, J.W. (2018). Integrins promote axonal regeneration after injury of the nervous system. Biol Rev 93, 1339–1362.

Ohyama, T., Jovanic, T., Denisov, G., Dang, T.C., Hoffmann, D., Kerr, R.A., and Zlatic, M. (2013). High-throughput analysis of stimulus-evoked behaviors in Drosophila larva reveals multiple modality-specific escape strategies. PLoS One 8, e71706.

Pease-Raissi, S.E., Pazyra-Murphy, M.F., Li, Y., Wachter, F., Fukuda, Y., Fenstermacher, S.J., Barclay, L.A., Bird, G.H., Walensky, L.D., and Segal, R.A. (2017). Paclitaxel Reduces Axonal Bclw to Initiate IP3R1-Dependent Axon Degeneration. Neuron 96, 373–386 e376.

Peltier, A.C., and Russell, J.W. (2002). Recent advances in drug-induced neuropathies. Curr Opin Neurol 15, 633–638.

Plantman, S., Patarroyo, M., Fried, K., Domogatskaya, A., Tryggvason, K., Hammarberg, H., and Cullheim, S. (2008). Integrin-laminin interactions controlling neurite outgrowth from adult DRG neurons in vitro. Mol Cell Neurosci 39, 50–62.

Poe, A.R., Tang, L.F., Wang, B., Li, Y., Sapar, M.L., and Han, C. (2017). Dendritic space-filling requires a neuronal type-specific extracellular permissive signal in Drosophila. P Natl Acad Sci USA 114, E8062–E8071.

Polydefkis, M., Yiannoutsos, C.T., Cohen, B.A., Hollander, H., Schifitto, G., Clifford, D.B., Simpson, D.M., Katzenstein, D., Shriver, S., Hauer, P., et al. (2002). Reduced intraepidermal nerve fiber density in HIV-associated sensory neuropathy. Neurology 58, 115–119.

Reichling, D.B., and Levine, J.D. (2011). Pain and death: neurodegenerative disease mechanisms in the nociceptor. Ann Neurol 69, 13–21.

Roberts, M., Barry, S., Woods, A., van der Sluijs, P., and Norman, J. (2001). PDGF-regulated rab4-dependent recycling of alphavbeta3 integrin from early endosomes is necessary for cell adhesion and spreading. Curr Biol 11, 1392–1402.

Roberts, M.S., Woods, A.J., Dale, T.C., van der Sluijs, P., and Norman, J.C. (2004). Protein kinase B/Akt acts via glycogen synthase kinase 3 to regulate recycling of alpha v beta 3 and alpha 5 beta 1 integrins. Molecular and Cellular Biology 24, 1505–1515.

Rong, Y., McPhee, C.K., Deng, S., Huang, L., Chen, L., Liu, M., Tracy, K., Baehrecke, E.H., Yu, L., and Lenardo, M.J. (2011). Spinster is required for autophagic lysosome reformation and mTOR reactivation following starvation. Proc Natl Acad Sci U S A 108, 7826–7831.

Schindelin, J., Arganda-Carreras, I., Frise, E., Kaynig, V., Longair, M., Pietzsch, T., Preibisch, S., Rueden, C., Saalfeld, S., Schmid, B., et al. (2012). Fiji: an open-source platform for biological-image analysis. Nat Methods 9, 676–682.

Schmidt, R.E., Dorsey, D., Parvin, C.A., Beaudet, L.N., Plurad, S.B., and Roth, K.A. (1997). Dystrophic axonal swellings develop as a function of age and diabetes in human dorsal root ganglia. J Neuropathol Exp Neurol 56, 1028–1043.

Seretny, M., Currie, G.L., Sena, E.S., Ramnarine, S., Grant, R., MacLeod, M.R., Colvin, L.A., and Fallon, M. (2014). Incidence, prevalence, and predictors of chemotherapy-induced peripheral neuropathy: A systematic review and meta-analysis. Pain 155, 2461–2470.

Shah, A., Hoffman, E.M., Mauermann, M.L., Loprinzi, C.L., Windebank, A.J., Klein, C.J., and Staff, N.P. (2018). Incidence and disease burden of chemotherapy-induced peripheral neuropathy in a population-based cohort. J Neurol Neurosurg Psychiatry 89, 636–641.

Shemesh, O.A., and Spira, M.E. (2010). Paclitaxel induces axonal microtubules polar reconfiguration and impaired organelle transport: implications for the pathogenesis of paclitaxel-induced polyneuropathy. Acta Neuropathol 119, 235–248.

Siau, C., and Bennett, G.J. (2006). Dysregulation of cellular calcium homeostasis in chemotherapy-evoked painful peripheral neuropathy. Anesth Analg 102, 1485–1490.

Soba, P., Han, C., Zheng, Y., Perea, D., Miguel-Aliaga, I., Jan, L.Y., and Jan, Y.N. (2015). The Ret receptor regulates sensory neuron dendrite growth and integrin mediated adhesion. Elife 4.

Soba, P., Zhu, S., Emoto, K., Younger, S., Yang, S.J., Yu, H.H., Lee, T., Jan, L.Y., and Jan, Y.N. (2007). Drosophila sensory neurons require Dscam for dendritic self-avoidance and proper dendritic field organization. Neuron 54, 403–416.

Sonnichsen, B., De Renzis, S., Nielsen, E., Rietdorf, J., and Zerial, M. (2000). Distinct membrane domains on endosomes in the recycling pathway visualized by multicolor imaging of Rab4, Rab5, and Rab11. J Cell Biol 149, 901–914.

Tanner, K.D., Levine, J.D., and Topp, K.S. (1998). Microtubule disorientation and axonal swelling in unmyelinated sensory axons during vincristine-induced painful neuropathy in rat. Journal of Comparative Neurology 395, 481–492.

Tomaselli, K.J., Doherty, P., Emmett, C.J., Damsky, C.H., Walsh, F.S., and Reichardt, L.F. (1993). Expression of beta 1 integrins in sensory neurons of the dorsal root ganglion and their functions in neurite outgrowth on two laminin isoforms. J Neurosci 13, 4880–4888.

Tracey, W.D., Jr., Wilson, R.I., Laurent, G., and Benzer, S. (2003). painless, a Drosophila gene essential for nociception. Cell 113, 261–273.

Tseng, M.T., Hsieh, S.C., Shun, C.T., Lee, K.L., Pan, C.L., Lin, W.M., Lin, Y.H., Yu, C.L., and Hsieh, S.T. (2006). Skin denervation and cutaneous vasculitis in systemic lupus erythematosus. Brain 129, 977–985.

Vonderheit, A., and Helenius, A. (2005). Rab7 associates with early endosomes to mediate sorting and transport of Semliki forest virus to late endosomes. PLoS Biol 3, e233.

Wallquist, W., Zelano, J., Plantman, S., Kaufman, S.J., Cullheim, S., and Hammarberg, H. (2004). Dorsal root ganglion neurons up-regulate the expression of laminin-associated integrins after peripheral but not central axotomy. J Comp Neurol 480, 162–169.

Wozniak, K.M., Vornov, J.J., Wu, Y., Liu, Y., Carozzi, V.A., Rodriguez-Menendez, V., Ballarini, E., Alberti, P., Pozzi, E., Semperboni, S., et al. (2018). Peripheral Neuropathy Induced by Microtubule-Targeted Chemotherapies: Insights into Acute Injury and Long-term Recovery. Cancer Res 78, 817–829.

Xiao, W.H., Zheng, H., Zheng, F.Y., Nuydens, R., Meert, T.F., and Bennett, G.J. (2011). Mitochondrial Abnormality in Sensory, but Not Motor, Axons in Paclitaxel-Evoked Painful Peripheral Neuropathy in the Rat. Neuroscience 199, 461–469.

York, A.G., Chandris, P., Nogare, D.D., Head, J., Wawrzusin, P., Fischer, R.S., Chitnis, A., and Shroff, H. (2013). Instant super-resolution imaging in live cells and embryos via analog image processing. Nat Methods 10, 1122–1126.

