## Supplemental Table 2 for "Integrins protect nociceptive neurons in models of paclitaxel-mediated peripheral sensory neuropathy"

Supplementary Table 2. List of genotypes used in this study

| Figure 1A-C | w-; ppk1.9Gal4/+; ppkcd4tdgfp/+ | Crossed with w^1118^ |
| --- | --- | --- |
| Figure 2A, C, C’ | w-; ppk1.9Gal4/+; ppkcd4tdgfp/+ | Crossed with w^1118^ |
| Figure 2B, D, D’ | w-; ppk1.9Gal4/+; ppkcd4tdgfp/ UASαPS1, UASβPS |  |
| Figure 2G | w^1118^ isogenized |  |
| Figure 2H | w-; ppk1.9Gal4/+; ppkcd4tdgfp/+  w-; ppk1.9Gal4/+; ppkcd4tdgfp/ UASαPS1, UASβPS | Crossed with w^1118^ |
| Figure 3A | w-; ppk1.9Gal4/UAS Rab4-mRFP; ppkcd4tdgfp/+ | UAS Rab4-mRFP is originated from BL8505 |
| Figure 3B | w-; ppk1.9Gal4/+; ppkcd4tdgfp/UAS Spin-myc-mRFP | UAS Spin-myc-mRFP is isolated from BL42716 |
| Figure 3H, I | w-; ppk1.9Gal4/UAS YFP-Rab4 WT; ppkcd4tdgfp/ UASαPS1, UASβPS | UAS YFP-Rab4 WT is originated from BL23269 |
| Figure 4A-D | Wildtype C57Bl/6J mice |  |
| Supp Fig 1 | w-; ppk1.9Gal4/+; ppkcd4tdgfp/+ | Crossed with w^1118^ |
| Supp Fig 2 | w-; ppk1.9Gal4/+; ppkcd4tdgfp/+ | Crossed with w^1118^ |
| Supp Fig 3 | w-; ppk1.9Gal4/ UAS-Ncad^7b-13a-18b^; ppkcd4tdgfp/+ |  |
| Supp Fig 4 | w-; UAS TrpA1/+; 412Gal4/+ | UAS TrpA1 is isolated from BL26263 |
| Supp Fig 5 | w-; ppk1.9Gal4/+; ppkcd4tdgfp/+  w-; ppk1.9Gal4/UAS βPS RNAi; ppkcd4tdgfp/+  w-; ppk1.9Gal4/+; ppkcd4tdgfp/ UASαPS1, UASβPS  w-; ppk1.9Gal4/ UAS-Ncad^7b-13a-18b^; ppkcd4tdgfp/+ | Crossed with w^1118^ |

Supp fig 5 ppk1.9Gal4, ppkcd4tdgfp were crossed with 1) w^1118^ (genetic control), 2) UAS Integrin β PS1 RNAi, 3) UASαPS1, UASβPS or 4) UAS Ncad, and
